# Expectation Violations as an Effective Alternative to Complex Mentalizing in Novel Communication

**DOI:** 10.1101/2024.11.19.624256

**Authors:** Tatia Buidze, Yuan-Wei Yao, Jan Gläscher

## Abstract

Effective communication in the absence of a shared language is a fundamental challenge, often addressed through complex cognitive mechanisms such as Theory of Mind, which allows individuals to infer others’ intentions and beliefs. However, this process is cognitively demanding and may not always be necessary. In this study, we propose that a more parsimonious cognitive mechanism—expectancy violations—can serve as an efficient alternative for communication in novel interactions. We tested this in the Tacit Communication Game, where we simulated Sender behavior using four computational models: the Surprise model based on expectancy violations and three levels of Theory of mind. After human Receivers interacted with these simulated Senders, we assessed the effectiveness of communication by analyzing accuracy and reaction times. Our results revealed that Receivers paired with the Surprise model achieved accuracy rates comparable to those interacting with the most complex Theory of mind model and exhibited more human-like message patterns. Additionally, models associated with higher accuracies also resulted in faster reaction times, indicating a reduced cognitive load. These findings challenge the necessity of complex mentalizing strategies in novel human interactions and suggest that an intuitive mechanism of expectancy violation may be a more plausible cognitive mechanism, while also providing quick responses.

## Introduction

Human communication often appears straightforward when viewed through the lens of classical theories like Shannon’s model^1^, where the process is simplified as encoding, transmitting, and decoding information through a physical medium with possibly signal degradation through interference by noise. However, this oversimplification leaves the challenges of an understanding of the meaning of a message unaddressed, especially when communicative conventions are absent or not shared. While established communicative systems, such as spoken language or gestures like raising a hand in a classroom, facilitate smooth communication by setting clear expectations, real-world scenarios sometimes lack such shared frameworks^2^. In these instances, communication becomes significantly more complex, requiring more than just the basic transmission of signals; it demands the use of recipient design^3,4^. This involves tailoring messages based on the Receiver’s likely perspective and understanding. This process often depends on Theory of Mind (ToM), wherein the Sender anticipates how the Receiver will interpret the signals based on their assumed mental state and knowledge^5–7^. While this perspective-taking mechanism is traditionally recognized as crucial for effective communication^8–11^, it can be cognitively demanding due to the need for accurate and detailed predictions about the Receiver’s state. Some argue that such cognitive effort may not be necessary in all communicative situations. Instead, straightforward strategies may be sufficient, allowing communicators to bypass detailed mentalizing by exploiting environmental cues or pre-established patterns^12–14^.

In this study, we examine a communicative scenario where the Sender must convey a goal location to the Receiver without using language or gestural cues, but solely through the movement trajectory of her token on a game board. This unique setting allows us to explore an alternative approach to the signaling function of communication: the expectancy violation mechanism. Instead of relying on complex ToM processes, this mechanism proposes that the Sender leverages violations of the Receiver’s fundamental expectations about the Sender’s movements to attract attention and signal crucial goal information^15^.

This approach is based on two key assumptions. First, it assumes that Receivers have fundamental expectations about the Sender’s movement, rooted in basic physical laws—such as the expectation that a moving object will continue along a straight path unless acted upon by an external force^16^. By intentionally deviating from these universally understood motion patterns, for example, by making unexpected turns, the Sender creates violations of these expectations to convey specific intentions. Second, the model assumes that the Sender expects the Receiver to detect these surprising events. While this may seem trivial, it is a crucial consideration because the effectiveness of expectancy violations depends on the Receiver’s ability to notice and interpret deviations from the norm. Organisms inherently need to detect changes in their environment to adapt to new conditions. Such changes are surprising because they represent discontinuities from previous environmental states, prompting attention and potential behavioral adjustments.

This approach resonates with the perspective that spatial cognition serves as a core organizational tool in human thought and that spatial concepts are universal^17^. Such cognition is fundamentally centered on the perceiver’s body^18–21^, and learning spatial expressions in a language largely involves mapping these universal concepts onto specific linguistic forms^22–24^. Our model incorporates these universal spatial priors by focusing on movement kinetics as a shared cognitive foundation. In this framework, expectancy violations leverage basic expectations about motion to create intuitive signals, effectively guiding attention without complex mentalizing.

The central question of this paper is twofold: Which approach is more effective in communicating a goal—the expectancy violation model or the more cognitively demanding ToM-based models— and which of these models more closely resembles human-generated messages? To investigate this, we use the Tacit Communication Game (TCG), which places participants in scenarios requiring non-verbal communication without pre-established conventions^25–27^. In this game, both players must reach their individual (non-identical) goal positions on a grid-like game board. However, only one of the players, the Sender (she/her) knows both goal locations, whereas the other player, the Receiver (he/his), only sees the Sender’s goal. The Sender then designs a movement trajectory (the “message”) and moves her token across the game board in such a way that communicates the goal position of the Receiver to him. In the previous study, we let the human Senders and Receivers play the game together and analyzed the human-generated movement trajectories. We observed that Senders successfully guided Receivers to their goal locations, effectively using movement trajectories that incorporated patterns like momentary diversions or zigzag movement^15^.

Building on these findings, we developed two computational approaches for this behavior: ToM-based models and the Surprise model. The ToM-based models incorporate varying levels of Theory of Mind, ranging from basic (ToM-0: no mentalization) to advanced (ToM-2: a recursive understanding that others hold beliefs about one’s own beliefs). In the TCG, the ToM-0 model assumes that the Sender selects paths without considering the Receiver’s mental state, relying purely on the physical characteristics of the path. The ToM-1 model introduces a more refined approach, where the Sender anticipates the Receiver’s likely interpretations of the path, aiming to minimize ambiguity by simplifying the route. This model captures a basic level of mentalizing, as the Sender adjusts message selection based on her awareness of the Receiver’s perspective. The most advanced ToM-2 model involves recursive mentalizing, where the Sender considers how the Receiver might infer the Sender’s intentions and adjusts the path accordingly to ensure clarity. These models span varying levels of cognitive complexity, providing a basis for comparison with the more parsimonious mechanism of expectancy violation.

In contrast, Surprise model leverages fundamental principles of movement to create communicative signals. We developed two types of priors: one based on movement kinetics— where it is expected that a moving object continues along its path unless an external force intervenes—and another based on the Sender’s goal orientation. These priors form a common language in the game, where straight-line trajectories align with expected directions. Deviations from these trajectories are used intentionally to create ‘surprise,’ drawing the Receiver’s attention and transforming these deviations into communicative acts. The Surprise model thus constructs messages that maximize information-theoretic surprise, turning expectancy violations into meaningful social signals.

In this study, we aimed to determine whether messages from the ToM and Surprise model are similar and equally well understood by the Receiver. To address this, we simulated the Sender’s behavior according to each model and had human Receivers interact with these simulated Senders, recording their accuracy and reaction times in identifying their goal states. We hypothesized that the expectancy violation mechanism, based on universal physical laws of motion, would be as effective as the more complex ToM models in guiding the Receiver’s attention and improving communication accuracy.

Our results support this hypothesis, revealing that the Surprise model achieved both high accuracy and low reaction times, comparable to those of the ToM-2 model, while outperforming the ToM-0 and ToM-1 models in both measures. In addition, we show that messages generated by the Surprise model resemble those messages compiled by human Senders more closely than those generated by the ToM-2 model. These findings suggest that an expectancy violation mechanism is indeed sufficient for effective non-verbal communication. This intuitive strategy, rooted in the violation of universally understood physical principles, can successfully attract the Receiver’s attention and guide them to the correct conclusions, offering a robust alternative to the more cognitively demanding ToM processes.

## Results

In this study, we investigated the effectiveness of various cognitive strategies in non-verbal communication by analyzing how Senders convey information to Receivers without shared language in the Tacit Communication Game (TCG). We simulated the behavior of the Sender in TCG, by employing four computational models: three different levels of Theory of Mind (ToM-0, ToM-1, and ToM-2) and a Surprise model — each designed to represent different cognitive strategies (see details of computational models in the Methods). A total of 40 participants played as Receivers and interacted with these simulated agents to assess the effectiveness of the different approaches. Each block in Figure 1a corresponds to one of these computational models, with each model being applied across 30 trials in the TCG. We analyzed the accuracy and reaction times of human Receivers interacting with these simulated Senders and compared the message patterns generated by the computational models with those produced by human Senders.

**Figure 1.**
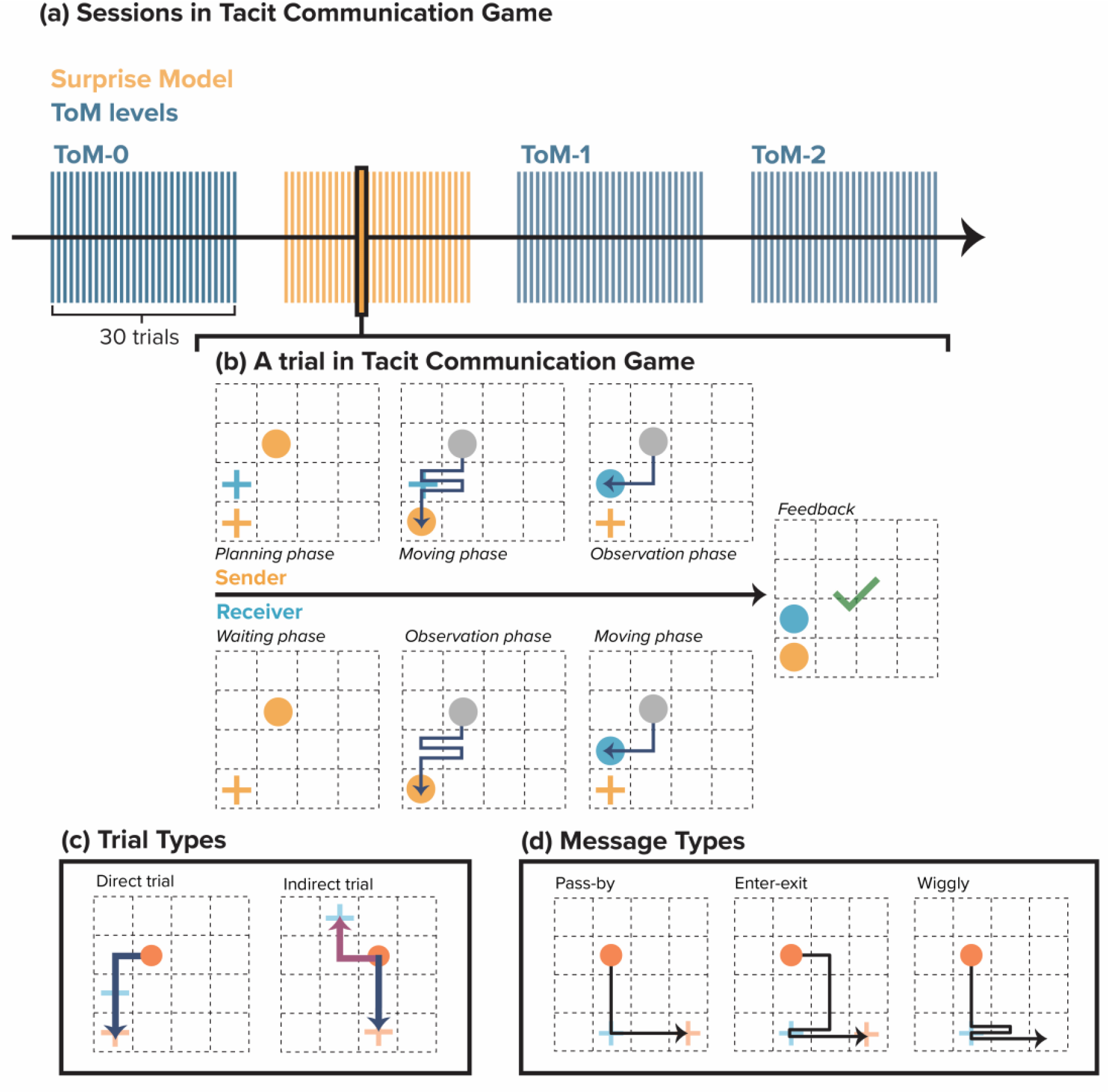
Experimental design of the Tacit Communication Game. (a) Sessions in Tacit Communication Game: Overview of the experimental design showing the four different sessions modeled after three levels of Theory of Mind — ToM-0, ToM-1, ToM-2—and the Surprise model, each consisting of 30 trials. (b) A trial in Tacit Communication Game: Phases of a trial in the Tacit Communication Game: (1) Planning phase where the Sender designs a path to communicate the Receiver’s hidden goal location, (2) Moving phase where the Sender executes the path, and (3) Observation phase followed by the Receiver’s movement and feedback. (c) Trial Types: demonstration of Direct and Indirect trial types used in the study: Direct trials require no deviation from the shortest path to the Sender’s goal, whereas indirect trials demand a strategic deviation to pass the Receiver’s goal. (d) Message Types: Types of messages used by the Sender to indicate the goal to the Receiver: Pass-by (simplest path), Enter-exit (explicit indication by entering and exiting the Receiver’s goal), and Wiggly (complex paths with multiple directional changes).

Below, we first provide a detailed description of the TCG, including the different goal configurations and example message trajectories. Next, we describe the computational models, focusing on the three levels of ToM and Surprise model. This is followed by an analysis of the message types generated by the different models and their influence on Receiver accuracy and reaction times.

### Tacit Communication Game

The Tacit Communication Game (TCG) is a cooperative, non-verbal task designed to investigate novel communication between a Sender and a Receiver without a shared language^11,25,26^. In this task, the Sender should create a movement path on a grid to communicate the hidden goal location of the Receiver (Figure 1b). The Receiver’s objective is to interpret the Sender’s movements and identify his own goal. The Sender has access to the starting positions and both goals—her own (Orange cross) and the Receiver’s (Blue cross)—whereas the Receiver is only aware of the starting location and the Sender’s goal. The Sender must plan a trajectory that communicates the Receiver’s goal and ends at her own goal, while minimizing the number of moves to maintain a high score. The score is quantified as 10 points reduced by the number of steps the Sender used to transmit the message. In this study, the Senders were simulated by computational models and the Receivers were human participants.

The trials were organized into two categories based on the spatial relationship between the goals: Direct and Indirect trials (Figure 1c). In Direct trials, the shortest path to the Sender’s goal passes by the Receiver’s goal, requiring no deviation. In contrast, Indirect trials require the Sender to deviate from a direct path to visit the Receiver’s goal before moving to their own. To convey these messages, the Sender can use different types of movement trajectories, which vary in their complexity and signaling strategies (Figure 1d). These include Pass-By messages, where the Sender stays on the direct path and briefly passes the Receiver’s goal; Enter-Exit messages, where the Sender explicitly diverges from the direct path to visit the Receiver’s goal; and Wiggly messages, characterized by zigzagging between the Receiver’s goal and the next state. These different strategies represent diverse communicative approaches and challenge the Receiver’s ability to find the correct goal location.

### Computational models

In this study, four computational models were used to simulate the behavior of the Sender in the TCG: the Surprise model and three Theory of Mind (ToM) models, which operate at varying levels of mentalizing complexity (ToM-0, ToM-1, and ToM-2). These models were tested to assess how well they conveyed information to human Receivers by analyzing the Receivers’ accuracies and reaction times.

The ToM models assume that the Sender selects messages based on their beliefs about the Receiver’s cognitive processes. The ToM models progressively increase in complexity, representing different levels of the Sender’s ability to consider the Receiver’s perspective when planning a path. In contrast, the Surprise model uses a different approach, focusing on expectancy violations to create communicative signals. We will briefly introduce both models below. For a more detailed description, please consult the Method section.

### Theory of Mind models

ToM models represent different levels of reasoning by the Sender in predicting how the Receiver interprets the observed trajectory. As the ToM level increases, the Sender becomes more sophisticated in guiding the Receiver toward the correct goal location by selecting messages that increasingly narrow down the possible number of Receiver goal states. We will briefly discuss the message selection and goal selection strategies on different levels below. For more details on how different levels of reasoning influence message types and their advantages across trial types, refer to the Methods section.

#### ToM-0 Model

The ToM-0 model assumes that the Sender has no awareness of the Receiver’s mental state and simply selects a movement pattern that minimizes the number of steps to retain the highest number of points. The Sender thus acts as a purely reward-maximizing agent with the primary objective of reaching her own goal while passing through the Receiver’s goal location. The ToM-0 Sender does not attempt to make the communication clearer for the Receiver beyond what is required to satisfy the basic rules of the game. For instance, in Figure 2a, the ToM-0 Sender would select the message on the left. The ToM-0 Receiver assumes that their goal is located on any tile along the Sender’s path, except the Sender’s final goal position, and selects a goal randomly from these options.

**Figure 2.**
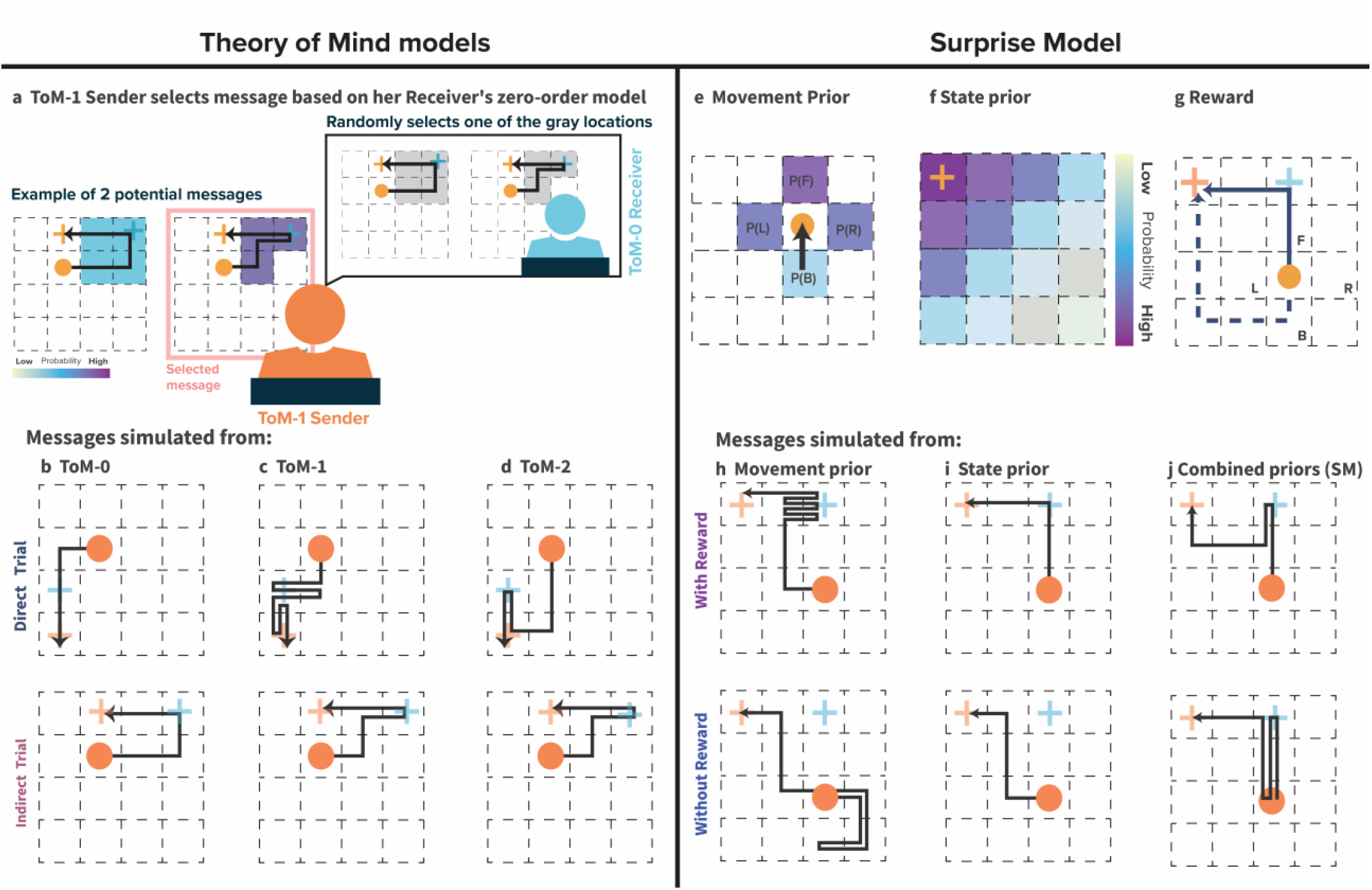
Computational models. (a) ToM-1 Sender selects a message based on the Receiver’s zero-order model: The ToM-1 Sender considers how the ToM-0 Receiver might interpret the path and selects the message that minimizes ambiguity. In this example, the Sender anticipates the Receiver’s random selection of a goal from the remaining path locations after excluding the Sender’s goal. This leads to a selection that aims to simplify the Receiver’s decision-making process by reducing the number of possibilities. (b) ToM-0 model generates Pass-by messages for both trial types, focusing only on efficiency without considering the Receiver’s perspective. (c) ToM-1 model creates Wiggly messages for direct trials and Enter-exit messages for indirect trials, using communicative signals to guide the Receiver. (d) ToM-2 model produces optimal Enter-exit messages for both trial types. (e) The Surprise Model’s Movement Prior assigns the highest probability to forward movements, followed by left and right, and the least for backward. (f) State Prior decreases as distance from the Sender’s goal increases (g) The Reward structure encourages efficient paths by rewarding actions that move towards the goal. (h) Movement prior emphasizes unexpected events to highlight the Receiver’s goal, while (i) State prior focuses on efficient Sender’s goal achievement. (j) The Surprise model combines both optimizing cost and signaling for the Receiver’s goal. When comparing messages with reward information (top row) to those without (bottom row), it becomes clear that including reward optimizes both cost and clarity in signaling the Receiver’s goal.

#### ToM-1 Model

The ToM-1 model introduces a higher level of reasoning, where the Sender attempts to consider how the ToM-0 Receiver might interpret the path (Figure 2a). The ToM-1 Sender tries to visit fewer grid states, since she knows that the ToM-0 Receiver will select the goal randomly among remaining locations on the message path after eliminating the Sender’s goal location (Figure 2a, right side message). This strategy helps the Receiver to infer his goal location more accurately by reducing the number of possibilities he needs to consider. The ToM-1 Receiver assumes that the ToM-0 Sender selects the shortest path and eliminates less likely goal locations based on simulating shorter alternative paths that the ToM-0 Sender could have sent but did not. By contrasting states of these shorter messages with the states in the observed message, the Receiver refines his possible goal locations.

#### ToM-2 Model

The ToM-2 model represents the most advanced level of mentalizing. Here, the Sender not only predicts the ToM-1 Receiver’s interpretation of the path but also anticipates how the Receiver might simulate alternative paths. The ToM-2 Sender intentionally chooses a message that deviates slightly from the most efficient path to ensure that the Receiver can easily detect the correct goal location. The ToM-2 Receiver uses a similar simulation strategy as the ToM-1 Receiver but refines their interpretation further by simulating multiple possible paths and eliminating unlikely options based on knowing that the ToM-1 Sender tries to minimize the number of states. This results in a more accurate selection of the goal location.

#### Different message types across levels of Theory of Mind and trial types

As the ToM models increase in complexity, we see different message types emerging depending on the Sender’s level of reasoning and the trial type. At the ToM-0 level, both direct and indirect trials typically result in what we previously described as a Pass-by message (Figure 2b). Since the ToM-0 Sender does not consider the Receiver’s perspective, the selected path is purely goal-oriented, without any additional communicative signals. The Sender simply follows the shortest route, leaving the Receiver to infer their goal by randomly selecting from possible options along the path.

At the ToM-1 level, the message types start to vary based on the trial type. In direct trials, the ToM-1 Sender generates Wiggly messages (Figure 2c, top), while in indirect trials, they produce Enter-exit messages (Figure 2c, bottom). These message types, unlike those in ToM-0, include intentional communicative signals. The ToM-1 Sender, having a basic understanding of how a ToM-0 Receiver interprets paths, tries to clarify the message by reducing possible goal locations, thereby guiding the Receiver more effectively. This shift indicates the Sender’s intention to communicate, anticipating the Receiver’s need for clearer cues.

At the ToM-2 level, the Sender’s communicative intention becomes even more pronounced. The ToM-2 Sender, predicting both the Receiver’s interpretation and potential path simulations, selects the most optimal and communicative Enter-exit messages for both direct and indirect trials (Figure 2d). These messages are designed to be easily distinguishable by the Receiver, providing the clearest signals for accurate goal identification. For more details on how different levels of reasoning influence message types and their advantages across trial types, refer to the Methods section.

### Surprise model

The Surprise model differs from the ToM models as it does not rely on recursive mentalizing. Instead, it is based on the principle of expectancy violations, where the Sender intentionally deviates from predicted movement patterns to capture the Receiver’s attention and signal important information.

The Surprise model consists of three core components: the Movement model, the State model, and the Reward^15^. The Movement model governs the likelihood of different actions based on movement priors derived from universal physical laws of object movement, namely that an object will continue moving in the same direction without an external force. Thus, the Movement model assigns the highest probability to forward movements and lower probabilities to lateral and backward movements (Figure 2e). The State model establishes a prior for each state on the grid, with higher probabilities assigned to states closer to the Sender’s goal (Figure 2f). The Reward component incentivizes the Sender to construct shorter, more efficient paths to their goal, while still incorporating surprising deviations to signal the Receiver’s goal (Figure 2g).

The Surprise is calculated by how much a chosen action deviates from the combined movement and state priors. Specifically, higher deviations from likely movement-state transitions lead to greater surprise, making these actions stand out more to the Receiver. In the first phase of message creation, the Sender selects actions that maximize this surprise, ensuring the Receiver’s goal is effectively communicated. After the Receiver’s goal is conveyed, the Sender shifts focus to efficiency, aiming to reach their own goal as directly as possible. This two-phase strategy enables the Surprise model to combine effective communication with movement efficiency.

Figure 2h-j demonstrates the influence of different model components on message planning. In simulations using only the Movement model (Fig. 2h, top), the Sender performs multiple zigzag movements due to the high surprise associated with backward actions. Conversely, the pure State model (Fig. 2i, top) generates a direct path message with no unexpected moves. When both models are combined in the Surprise model (Fig. 2j, top), the result is similar to the human-generated Enter-exit messages. The second row shows the impact of removing the reward. This results in a loss of motivation to reach the Sender’s goal quickly and creates shorter messages to retain a high number of points. Without it, both the Movement and State models (Fig. 2h, i, bottom) produce similar message patterns but fail to communicate the Receiver’s goal. The Surprise model (Fig. 2j, bottom) results in 2-step wiggles, but without the reward component, it struggles to efficiently guide the Sender to their final goal.

In summary, combining Movement and State priors with a surprise action policy and future rewards allows the Sender to clearly signal the Receiver’s goal. Once this is achieved, the State priors then direct the Sender along the shortest path to their own goal.

### Message types generated by different models and their accuracies

In analyzing the computational strategies within the Tacit Communication Game, we identified three key message types—Pass-By (PB), Enter-Exit (EE), and Wiggly (W)—each reflecting a unique approach to non-verbal signaling by the Sender (Figure 1d).

#### Pass-By (PB) Messages

Most frequently utilized by the ToM-0 model, these messages constituted 87% of its communications (Figure 3b, upper row). Due to their straightforward and less informative strategy, PB messages generally yielded lower accuracy, as shown in Figure 3b (lower row).

**Figure 3.**
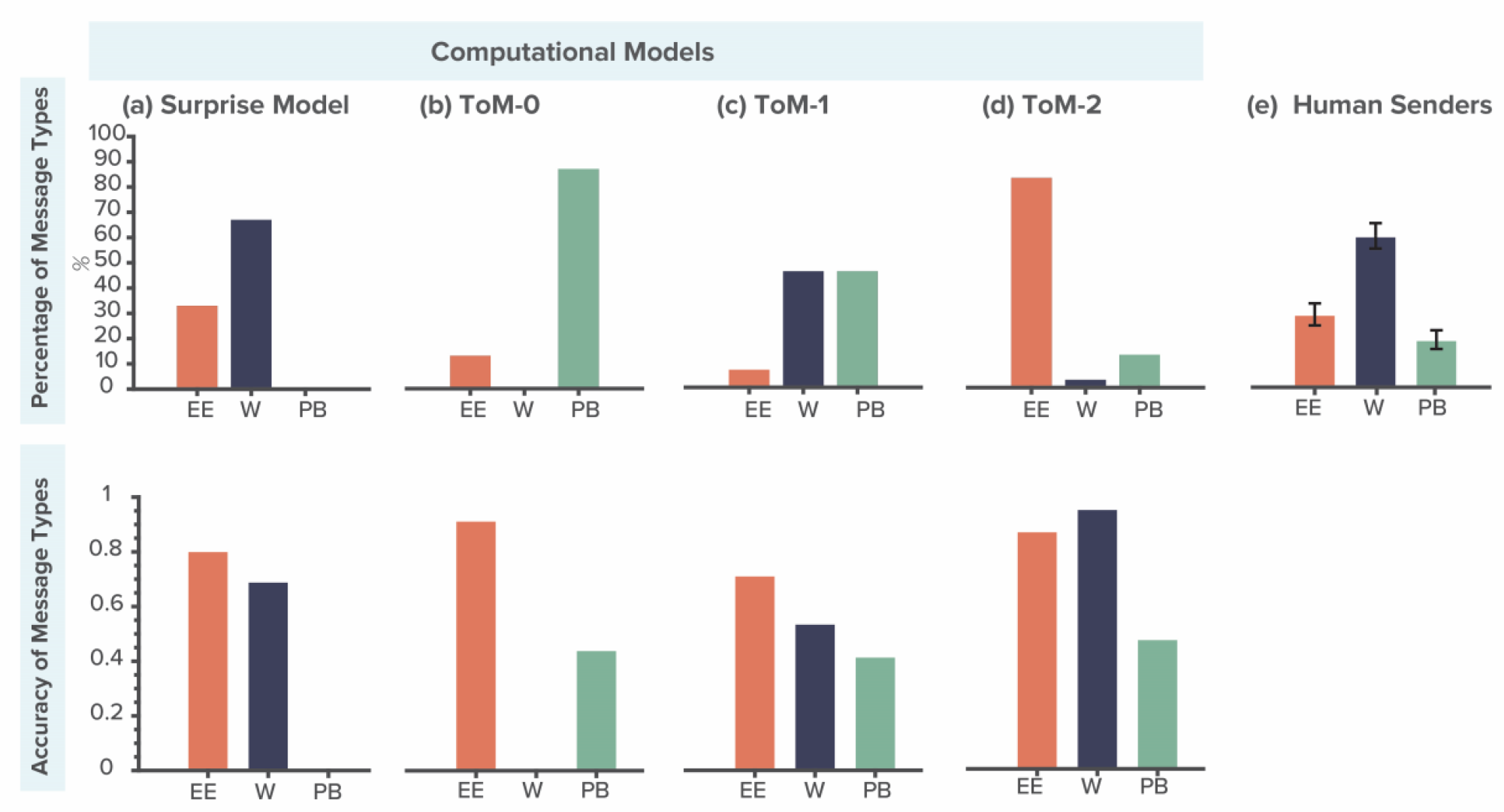
Analysis of message types across computational models and comparison with human Senders. (a) Surprise Model: Displays the distribution of message types generated by the Surprise model, with a predominant use of Wiggly (W) messages shown in dark blue, followed by Enter-Exit (EE) messages in orange. The accuracy graph below indicates that EE messages achieve the highest interpretation accuracy, followed by W. (b) ToM-0: Illustrates that the ToM-0 model predominantly uses PB messages. While EE messages are less frequently used, they maintain high accuracy. (c) ToM-1: Shows a more balanced distribution between EE and W messages, with the highest EE message accuracy followed by the W. PB messages are less frequent and exhibit lower accuracy compared to EE and W messages. (d) ToM-2: Presents the ToM-2 model, where W messages display higher accuracy than EE messages. However, W message frequency is very low compared to EE messages. (e) Human Senders: Compares the message type usage of the Surprise model with human Senders. Data are taken from previous study^15^. The graph highlights that human Sender’s frequencies of EE and W messages closely resemble the pattern observed in the Surprise model

#### Enter-Exit (EE) Messages

Heavily favored by the ToM-2 model, EE messages accounted for 83% of its communications (Figure 3d, upper row) and demonstrated high accuracy due to their explicit and direct signaling, as indicated in Figure 3d (lower row). The Surprise model also employed EE messages, representing 33% of its communications (Figure 3a, upper row), showing a preference for a direct approach with similarly high accuracy.

#### Wiggly (W) Messages

Employed by the ToM-1 model for 46% of its communications (Figure 3c, upper row), Wiggly messages suggested a moderate level of strategic complexity with corresponding moderate accuracy, as detailed in Figure 3c (lower row). The Surprise model, which used this message type in 66% of instances (Figure 3a, upper row), balanced complexity and effectiveness in signaling with moderate accuracy.

EE messages were consistently the most accurately interpreted by Receivers, irrespective of the model used, followed by W and then PB messages (Figure 3a-d, lower row). The only exception was in the ToM-2 model, where Wiggly messages showed higher accuracy than Enter-Exit (figure 3d, lower row). However, this is likely due to the lower frequency of Wiggly messages for this model (figure 3d, upper row), which could lead to an overestimation of its effectiveness.

Overall, these findings show that both the Surprise and ToM-2 model generated highly effective message types. However, comparative analysis with human Sender profiles (Figure 3e) revealed that the Surprise model closely resembles human-like messaging ^15^. This similarity suggests that the Surprise model is not only parsimonious and effective, but also reflects the strategies humans naturally use in non-verbal communication.

### Receiver’s accuracy and reaction times to different models

Analyzing the human Receiver’s accuracies using a One-way ANOVA analysis revealed significant differences among the tested models in communicating the Receiver’s goal location effectively (*F (3, 156) = 22*.*22, p* < *0*.*001, η*^*²*^ *= 0*.*3*). Specifically, the Surprise model (SM) and the ToM-2 model demonstrated the highest mean accuracies among the tested models (Figure 4a), with mean accuracies of *0*.*67* (SM) and *0*.*78* (ToM-2), respectively. Post-hoc comparisons using Tukey’s HSD test revealed significant differences between the Surprise model (SM) and both the ToM-0 and ToM-1 models. Specifically, the SM exhibited significant differences compared to ToM-0 (T = 2.5, *p* < *0*.*001*) and ToM-1 (*T = 2*.*84, p* < *0*.*001*), but did not differ significantly from the ToM-2 model (*T = 7*.*8, p > 0*.*001*). Similarly, the ToM-2 model exhibited comparable trends, with significant disparities observed between it and ToM-0 (*T = 14*.*3, p* < *0*.*001*), as well as ToM-1 (*T = 14*.*65, p* < *0*.*001*). This finding suggests that both the utilization of surprise elements in the SM and the incorporation of sophisticated inferential thinking in the ToM-2 performs significantly better than the ToM-0 and ToM-1 models, and that they both are similarly effective in facilitating the Receiver’s understanding of the game.

**Figure 4.**
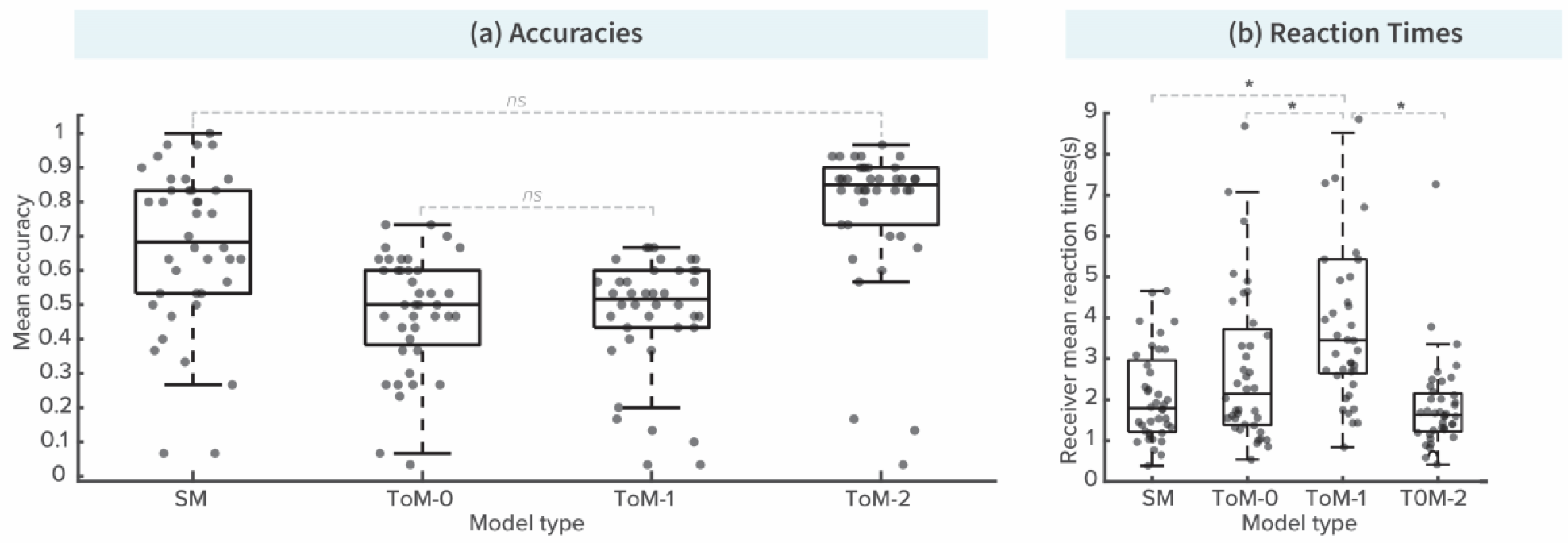
Performance metrics of different computational models in the Tacit Communication Game. (a) Accuracies: The graph displays the mean accuracy of each model in the game, with the Surprise Model (SM) and ToM-2 showing the highest accuracy. These models significantly outperformed the ToM-0 and ToM-1 models. No significant difference was found between the SM and ToM-2, indicating their comparable effectiveness. (b) Reaction Times: This graph illustrates the Receiver’s mean reaction times across different models. Faster reaction times were noted for the SM and ToM-2. Significant differences in reaction times were observed between the lower-level and higher-level ToM models, highlighting the impact of model complexity on processing speed.

Subsequent analysis revealed the influence of model behavior on the Receiver’s reaction time (*F (3,80) = 10*.*4, p = p* < *0*.*001, η*^*²*^ *= 0*.*2*). Models with higher accuracy, namely SM and ToM-2, demonstrated shorter reaction times for Receivers (Figure 4b). Specifically, average reaction times were shorter for SM (2.26 seconds) and ToM-2 (1.83 seconds) compared to those of ToM-0 (2.96 seconds) and ToM-1 (4.3 seconds). Notably, three significant differences emerged in post hoc Tukey’s HSD tests: between ToM-1 and SM (*T = 10*.*69, p* < *0*.*001*), ToM-0 and ToM-1 (*T = 8*.*40, p = 0*.*02*), and between ToM-1 and ToM-2 (*T = 4*.*12, p* < *0*.*001*). This link between high accuracy models and reduced reaction time may reflect a lower cognitive load for the Receiver due to the effective messaging during the communication process.

## Discussion

In our study, we investigate the cognitive processes underlying novel human interactions by employing the Tacit Communication Game (TCG) as a controlled experimental setup. We explored whether effective communication in this context is better facilitated by intuitive surprises or sophisticated mentalizing strategies. To this end, we compared the Surprise Model (SM), which leverages unexpected deviations from expected behavior, against three levels of recursive Theory of Mind (ToM). These ranged from ToM-0 where the Sender does not attempt to guess the Receiver’s thoughts, to ToM-2, where the Sender anticipates the Receiver’s interpretation of the message. Our results demonstrate compelling evidence that SM is similarly effective as the more complex ToM processes, particularly ToM-2, in enhancing understanding and ensuring efficient communication in the absence of shared language. Both the SM and ToM-2 achieved high accuracy rates and were linked to reduced reaction times for Receivers in decoding messages, suggesting that both models are equally viable for improving non-verbal communication in TCG.

A fundamental aspect of the SM is its proposition that unexpected behavior plays a crucial role in signaling intentions during communication, particularly when linguistic or cultural cues are absent. In these situations, individuals tend to revert to fundamental principles that inform human interaction—principles that are universally recognizable. These shared expectations form a basis upon which new forms of communication can develop through repeated social exchanges. By leveraging these common understandings, communicators can use surprising actions to effectively capture attention and highlight essential information. This approach diverges sharply from traditional communication models that rely on word predictability to facilitate comprehension^28,29^; instead, it embraces unexpectedness to enhance understanding in challenging contexts.

For example, the Rational Speech Act model posits that speakers optimize their utterances to minimize the listener’s surprise while maximizing the likelihood of accurate interpretation^28^. Similarly, the Information Density Hypothesis suggests that speakers manipulate syntactic structures to ensure a consistent flow of information, aligning their linguistic choices with the statistical regularities of speech^29^. Concepts from information theory, such as KL divergence or Shannon surprise, can also be utilized to assess word predictability within sentences^1^. While both the SM and conventional models engage with Shannon surprise, they do so with different intentions. Traditional models seek to minimize surprise for improved clarity, whereas the SM strategically maximizes it to convey information more effectively in settings devoid of shared language. This emphasis on offsetting expectations becomes a vital tool for communication in such contexts.

Several lines of research support the efficacy of communication through surprising events. Unexpected actions are often interpreted as intentional, allowing for efficient transmission of meaning^30^. Additional studies indicate that context can significantly influence selection^31^; for instance, participants often preferred less typical items when color was critical for identification— opting for a yellow chair rather than a banana. This adaptability illustrates how humans modify their expectations based on specific communicative situations, aligning well with the principles of SM.

A key advantage of the SM over the ToM-2 lies in its fundamental simplicity. By operating on fewer assumptions, the SM utilizes universal principles of expectancy violations, making it inherently less complex and more robust. Simpler models typically have fewer points of failure, enhancing their reliability^32,33^. The SM’s straightforward reliance on expectancy violations reduces the risk of misinterpretation that can occur with more complex models, which often require a certain level of cognitive capability in the Receiver. This clarity and reliability are particularly valuable in different communication environments where the predictability of responses might not be guaranteed. These characteristics make SM an efficient tool across a wide range of contexts, maintaining its effectiveness without the need for complex mentalizing processes. Moreover, the simplicity of the SM may facilitate its broader adoption in real-world applications and its generalization to other tasks more effectively than the ToM-2 models^34,35^.

An additional advantage of the SM over ToM models is its core operational mechanism. Unlike the ToM models that focus on message selection, the SM employs a step-by-step message creation process. ToM models rely on complex reasoning processes that involve defining a belief distribution over potential states on the game board and performing an exhaustive search for optimal messages for each goal configuration. This method, while theoretically sound, demands extensive cognitive resources and conflicts with the brain’s preference for cognitive efficiency. In contrast, the SM’s mechanism aligns more closely with biological plausibility, as it does not require the exhaustive cognitive load imposed by recursive mentalizing strategies. This aspect of the SM suggests that it may be more reflective of how the human brain naturally processes non-verbal communication, favoring strategies that balance efficiency with effectiveness.

As previously mentioned, the high accuracy rates observed with both the SM and the ToM-2 were accompanied by significantly quicker reaction times among Receivers. These shorter reaction times indicate that the information transmitted by the Sender was processed efficiently, enabling rapid decision-making. This efficiency is crucial in time-sensitive situations and underscores the efficacy of both communication strategies in reducing cognitive load during the communication process^36,37^.

Additionally, the SM generated messaging patterns that were more human-like. This resemblance to natural human communication suggests that the cognitive mechanisms employed by the SM closely mirror those instinctively used by humans in similar situations. This human-like messaging is crucial because it aligns with the natural tendencies of human communicators, offering greater ecological validity and potentially making interactions smoother and more intuitive in human-to-human contexts.

The Surprise model not only has simple and intuitive assumptions, but it also has innovative aspects. Unlike the typical ‘action selection by value’ approach seen in reinforcement learning^38^ and predictive coding^39^, the Surprise Model employs an “action selection by surprise” strategy. This unique method of decision-making focuses on disrupting established expectations to convey crucial information. Future research could explore how this approach can be integrated with AI technologies to enhance interaction between humans and machines, potentially through algorithms that prioritize unexpected choices to enhance engagement and creativity. For example, generative AI models trained on extensive text data typically analyze possible word sequences based on contextual information to ensure clarity and coherence, selecting the most probable words. While these models sometimes introduce variability through techniques like top-k sampling and temperature scaling, this only results in moderate diversity in word selection. By applying the principles of the Surprise model, AI could select significantly more unexpected words, thereby increasing variability in the generation process. This novel approach would diverge from predictable paths, enhancing creativity and engagement. Such an application could improve AI’s capabilities in areas like comedy, storytelling, or generative art, creating more engaging and unique scenarios.

While the findings are robust, this study is not without its limitations. The controlled nature of the experimental setup does not fully encapsulate the complexity of real-world communication scenarios, where environmental and interpersonal variables can significantly affect outcomes. It is important to note that while SM implements an effective signaling function of communication, this is just one characteristic of the broader communicative landscape. There are many aspects of communication where the straightforward SM may not be the optimal choice, particularly in contexts requiring more nuanced or hierarchical reasoning. These limitations highlight the need for caution when generalizing these findings to more complex, real-world interactions. Future research should examine the applicability of SM principles beyond the TCG and explore how the Surprise Model can be adapted or combined with other cognitive strategies to handle more complex communicative demands and encompass the broader facets of human communication.

In summary, our study demonstrates that the Surprise model, by utilizing a parsimonious and intuitive mechanism of expectancy violation, communicates as effectively as the more complex Theory of Mind processes. This effectiveness, combined with quicker reaction times and human-like messaging patterns, highlights its potential for broad application in enhancing non-verbal communication across different contexts and technologies.

## Methods

### Experimental design

The Tacit Communication Game (TCG) is a non-verbal cooperative task played by two players: a Sender (she/her) and a Receiver (he/his). The success of the game depends on the Sender’s ability to effectively construct a path on a grid that communicates the Receiver’s hidden destination, and on the Receiver’s capacity to accurately interpret this communicative message to find his goal location. Our study primarily delves into the Sender’s strategic behavior and the cognitive mechanisms that enable successful and effective non-verbal communication.

#### TCG sessions

To empirically test these theories, we simulated Sender behavior across three different levels of Theory of mind: ToM-0, ToM-1, and ToM-2—and contrasted them with a Sender following the Surprise model. This created 4 simulated Senders, each embodying a distinct communicative approach based on a particular cognitive model. Human Receivers were paired with each simulated Sender across four separate sessions (Figure 1a), with each session encompassing 30 trials. The trials represented different goal configurations, requiring the Sender to convey the target location through their movement across the grid. The sequence in which these sessions and their internal trials were presented was randomized to ensure variability and to mitigate any sequential learning effects.

#### TCG trial

Each trial began with a visual presentation on computer monitors. The Sender’s display included three key pieces of information: her own starting location depicted as an orange circle, and two goal locations marked by colored crosses— orange for the Sender and blue for the Receiver (Figure 1b, planning phase). In contrast, the Receiver’s display was limited to just the starting location and the Sender’s goal, with his own goal remaining undisclosed (Figure 1b, waiting phase). The central challenge of TCG emerges here, as the Sender must reach three objectives: to craft a trajectory that conveys the hidden goal to the Receiver, to ensure her path concludes at her goal position and to minimize the number of moves to maximize the score at the end of the trial. After the Sender’s move, the Receiver observes the trajectory (Figure 1b, Observation phase) and proceeds to deduce his target location based on the observation. He then navigates his token towards what he believes to be the correct destination (Figure 1b, Receiver’s movement phase). The trial culminates with feedback that reflects the effectiveness of communication. In each trial, players start with a score of 10 points, decreasing with each move the Sender makes, thus incentivizing the Sender to communicate the path in as few moves as possible.

#### Trial types

Trials within the TCG were divided into two categories based on the positional relation of the goals: Direct and Indirect (shown in Figure 1c). In Direct trials, the shortest path to the Sender’s goal, passes by the Receiver’s goal, demanding no deviation. In contrast, Indirect trials required the Sender to deviate from the direct path to visit the Receiver’s goal. We included 20 Direct trials and 10 Indirect trials in each session. We chose more direct trials because they are more challenging and have shown to elicit more diverse strategies in previous observations.

#### Message types

In examining non-verbal communication strategies within the Tacit Communication Game, we identified three primary message types—Pass-By, Enter-Exit, and Wiggly produced by different models at varying frequencies, with each type exemplifying a unique method of non-verbal signaling by the Sender (Figure 1d). The Pass-By Message involves Senders briefly passing through the Receiver’s goal location on their way to their own goal. The Enter-Exit Message involves Senders diverging from their direct route to the Receiver’s goal before proceeding to their own goal. Lastly, the Wiggly Message is characterized by Senders navigating on the direct path but shifting between the Receiver’s goal and the next state to create a zigzag pattern.

### Participants

We recruited 40 participants for the role of the Receiver in the Tacit Communication Game (TCG) using university job boards. We ensured that these individuals had no history of neurological or psychiatric diseases as part of our selection process. The study was approved by the General Medical Council of the City of Hamburg (Approval Code: PV7114). All participants were given detailed instructions about the study in written form and provided their written consent before participating. Participants were paid for their time, receiving a base rate of €10 per hour. They could also earn an extra bonus based on their performance, which averaged €4, to encourage their best effort during the experiment.

### Computational models

#### Theory of Mind model for Tacit Communication Game

The fundamental principle of employing Theory of Mind in a Tacit Communicative Game (TCG) relies on the assumption that the Sender designs a communicative signal based on her beliefs about the Receiver. She takes the Receiver’s perspective to predict his actions, and then designs a message that maximizes his chance of correctly identifying his goal location^11^. Similarly, the Receiver tries to put himself in the Sender’s position and guess for which goal location the Sender would use the observed signal^27^.

In the subsequent sections, we describe the computational ToM model and how this perspective-taking influences the strategies and behaviors of players at different levels of Theory of Mind within the context of the TCG. This interplay of mental models determines how effectively players can communicate and understand each other non-verbally, shaping the outcomes of the game. It is important to point out that all ToM models *select* messages that match the mentalizing operation of the ToM level from the set of all possible messages. In contrast, the Surprise model (described below) *constructs* messages step-by-step using a value iteration algorithm that capitalizes on the information-theoretic surprise of each possible next step.

### Definitions

#### Goal configuration and main message set

The goal configuration of each trial in the TCG is defined by 3 elements: a starting point *A*, the Sender’s goal location *G*_*s*_ and the Receiver goal location *G*_*r*_ . One of the building blocks of the ToM model of the TCG is the definition of a main message set *M*. This set consists of all the possible messages consisting of a number of states *s* that start at Sender’s goal location *A*, pass through the Receiver’s goal location *G*_*r*_ and end at Sender’s goal location *G*_*s*_. Formally we can define *M* as:

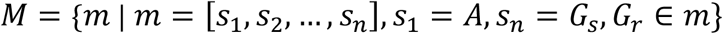

#### Distinction between message sequence and unique states

Message sequence *m* is a sequence of states that represent a path on the grid. Each state *s*_*i*_ corresponds to a specific position on the grid. We denote message sequence by *m =*[*s*_1_, *s*_2_, …, *s*_*n*_], where *s*_1_ *= A* is the Sender’s starting location and *s*_*n*_ *= G*_*s*_ is the Sender’s goal location. The length of a message sequence *L*( *m*) is the total number of the states in the sequence, given by *L*( *m*) *=* | *m*|. Within a message sequence, certain states may be revisited, resulting in fewer unique states than the sequence length. We define *U( m*) as the set of unique states in the message *m*:

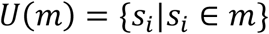

The number of unique states in *m* is then |*U( m*)|, which may be smaller than *L( m*) due to possible revisits.

#### General mechanism of creating level-specific message sets and goal selection

At each Theory of Mind (ToM) level *k* the Sender employs a unique strategy to create a message set, based on their mental model of a *k* − 1 level Receiver. This message set is crafted to help the Receiver narrow down potential candidate locations, thereby increasing the likelihood that the Receiver will correctly identify their goal location. The specific approach for constructing these sets varies by ToM level, which determines the types of messages included. The Receiver’s task, in turn, is to identify their own goal location. Like the Sender, the Receiver generates sets of possible goal locations, with the elements in these sets varying according to the Receiver’s ToM level. After forming a level-specific set, the Receiver randomly selects a goal location from the identified candidate locations.

In the following sections, we utilize the definitions above to explore this general mechanism, detailing the distinct message sets and goal selection strategies across ToM levels for both Sender and Receiver.

### ToM-0 players

#### Sender

##### Mental model

The ToM-0 Sender has no mental model of the Receiver but knows the rules of the game and wants to be successful. The Sender must meet three primary objectives of the game; the first is to reach her goal location; the second is to communicate the Receiver’s goal; the third is to make as few steps as possible.

##### Message selection

With all these objectives in mind, the ToM-0 Sender selects a movement pattern that passes through the Receiver’s goal location and ends in her goal location. As each step incurs a cost of 1 point, the Sender tries to minimize the length of the message. Thus, the ToM-0 Sender model defines the set of possible messages *M*_0_ (subscript indicating ToM level) as

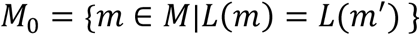

This means that each message in *M*_0_ has the minimum number of states in its sequence. Once the message set is created then the Sender randomly selects one message from *M*_0_:

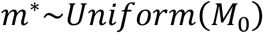

Meaning:

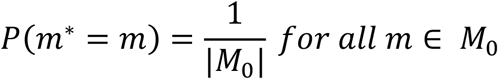

Figure 5a shows two possible messages, and the ToM-0 Sender selects Message 2 over Message 1 because it is shorter, thus conserving more points.

**Figure 5.**
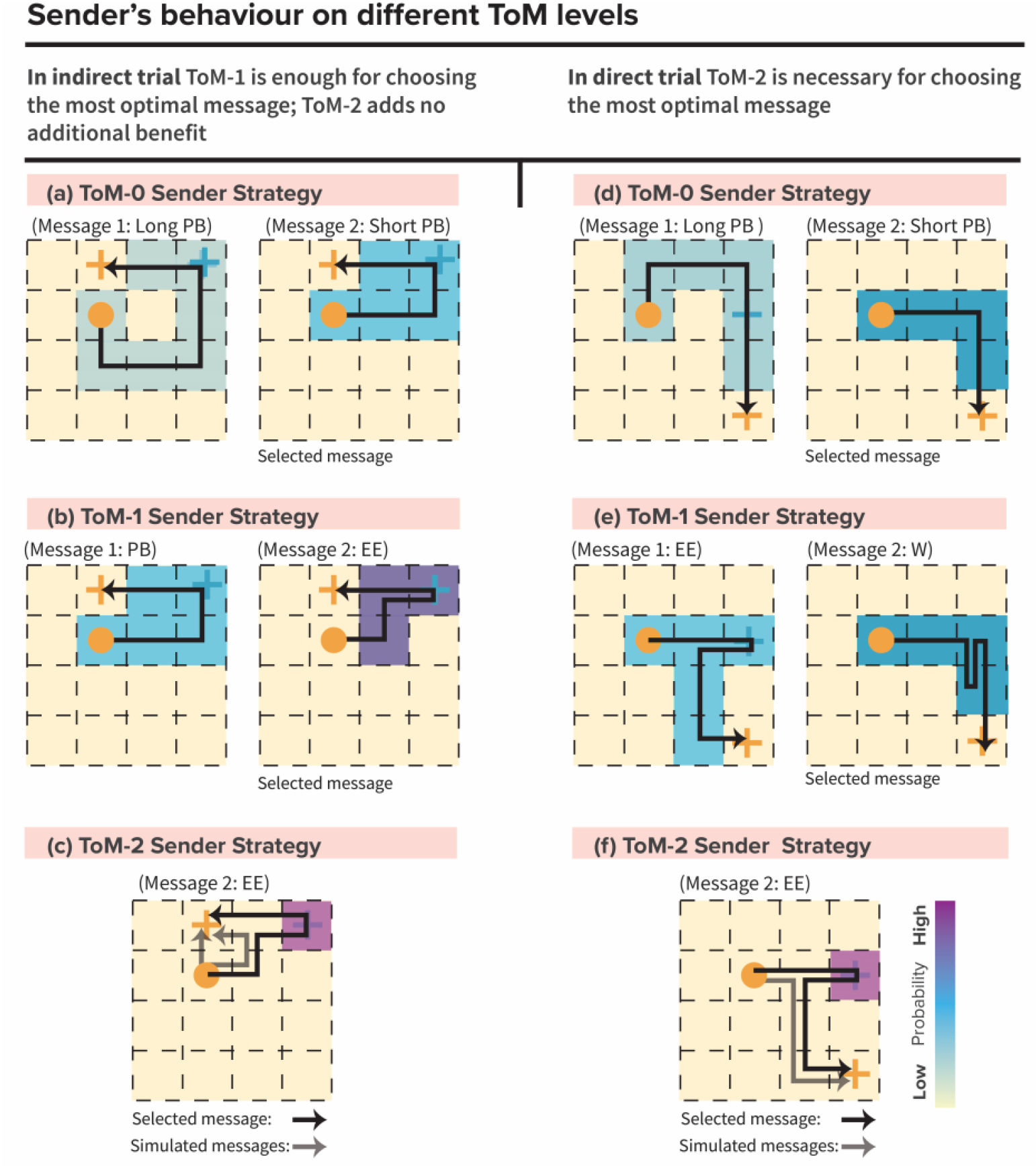
Sender’s behavior on different ToM levels. (a) ToM-0 Sender Strategy in Indirect Trial: For ToM-0, optimal message selection is simple; it selects the shortest path that passes by the Receiver’s goal. The example shows that Message 2 is selected for its shorter path. (b) ToM-1 Sender Strategy in Indirect Trial: ToM-1 strategy involves predicting the Receiver’s interpretation of the path, leading to a choice of a path with fewer visited states. The example shows the choice between two paths, selecting the one that minimizes state visits (message 2). (c) ToM-2 Sender Strategy in Direct Trial: ToM-2 explicitly deviates from the shortest route to ensure the Receiver can distinguish the intended message from simpler alternatives. (d) ToM-0 Sender Strategy in Direct Trial: In direct trials, the ToM-0 strategy still focuses on the shortest path without extra deviations, simplifying the Receiver’s decoding process. (e) ToM-1 Sender Strategy in Direct Trial: ToM-1 adjusts its path based on the Receiver’s expected mental reasoning, leading to selection of wiggly message (message 2). (f) ToM-2 Sender Strategy in Direct Trial: ToM-2 Sender, similar to their approach in indirect trials, intentionally deviates from the direct path.

### Receiver

#### Mental model

Similarly to the ToM-0 Sender, the ToM-0 Receiver has no mental model of his partner but knows about the rules of the game and wants to be successful.

#### Goal selection

The ToM-0 Receiver observes the Sender’s movement pattern and assumes that his goal location can be on any given state on the observed message path *m*^*^=[*s*_1_, *s*_2_, …, *s*_*n*_], except for the starting point *A* and the Sender’s goal *G*_*s*_. The model for the ToM-0 Receiver therefore constructs the following set of possible states for the Receiver’s goal location:

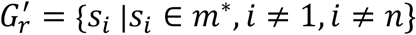

The Receiver randomly selects one state from 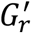:

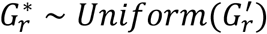

Figure 6b shows candidate locations depicted by blue color after seeing the message displayed in figure 6a. The Receiver will randomly select from these 4 goal locations.

**Figure 6.**
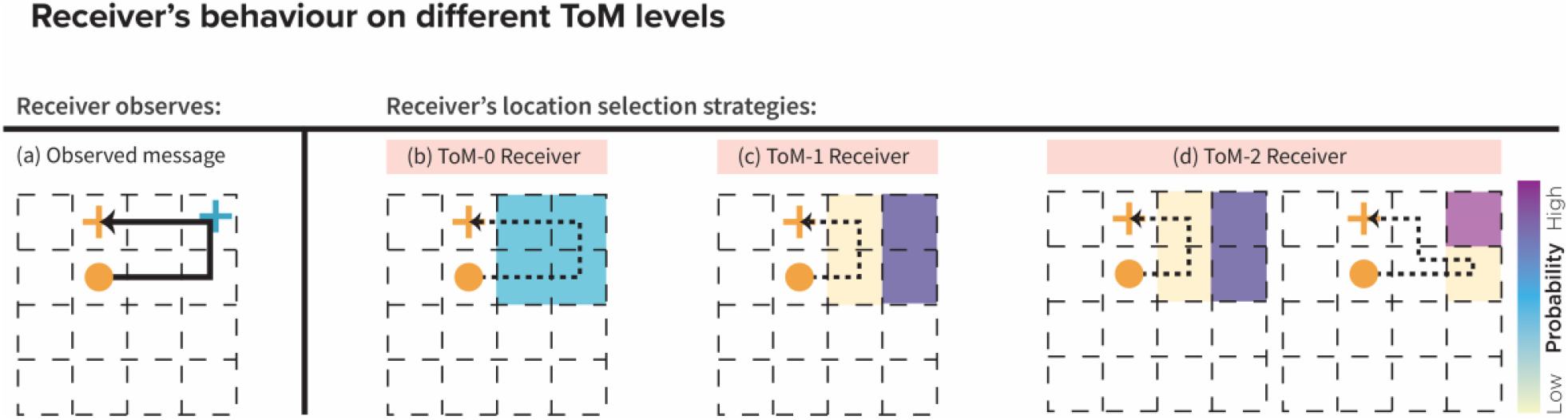
Receiver’s behavior on different ToM levels. (a) Observed Message: Illustration of the Sender’s path used in communication. The path depicted in black indicates the movement, with the Sender’s and Receiver’s goal locations marked by an orange circle and blue cross, respectively. (b) ToM-0 Receiver: ToM-0 Receiver assumes the goal could be any tile on the Sender’s path. The blue shaded tiles represent the Receiver’s potential goal locations after excluding the Sender’s end position. This Receiver selects his goal randomly from these highlighted locations. (c) ToM-1 Receiver: The ToM-1 Receiver employs a slightly more strategic approach, eliminating unlikely goal locations based on a shorter path logic. Yellow shaded tiles depict eliminated options, while the remaining purple tiles are considered equally probable as the goal location. (d) ToM-2 Receiver: ToM-2 Receiver further refines the elimination process, simulating potential shorter paths the ToM-1 Sender might have taken. The left grid shows initial elimination (yellow tiles), and the right grid shows further narrowed choices. The pink tile indicates the final selected goal location with high certainty, following the elimination of less probable options (yellow tiles on both grid).

### ToM-1 players Sender

### Mental model

ToM-1 Sender considers the game from the perspective of her ToM-0 partner. While designing the communicative signal, the ToM-1 Sender puts herself in the position of the ToM-0 Receiver and predicts that the Receiver will select the goal randomly among the remaining locations on the message path after eliminating the Sender’s goal location and the starting point.

#### Message selection

Unlike the ToM-0 Sender, who tries to make as few moves as possible, the ToM-1 Sender attempts to visit as few number of unique states as possible. To satisfy this constraint, the model constructs the following set of messages *M*_1_ as follows:

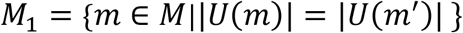

and selects one of them randomly. Formally:

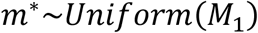

Meaning:

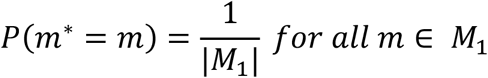

The difference between strategies used by the ToM-0 and ToM-1 Sender is shown in Figure 5b (message 1 and message 2). Even though both messages are the same length (5 steps), the ToM-1 message has only four visited states (depicted by purple-colored tiles on the grid), while the ToM-0 has five. Minimizing not only the length of the path, but also the number of unique states, the Sender has effectively increased the ToM-0 Receiver’s chance of correctly guessing his goal location from 0.2 to 0.25, because she knows that the ToM-0 Receiver is still selecting randomly from the set of possible message states, but which now contains one state less.

### Receiver

#### Mental model

ToM-1 Receiver has a ToM-0 model of the Sender, i.e., he knows that the ToM-0 Sender only selects the shortest message that passes through the Receiver’s and Sender’s goal locations.

#### Goal selection

The ToM-1 Receiver observes the Sender’s message *m*^*^ and aims to pinpoint his goal location by simulating less likely (shorter) messages *m*_*sim*_ and then calculating the difference between the two sets of states of the simulated and observed messages. Knowing that the ToM-0 Sender prefers the shortest path passing through his (the Receiver’s) goal, the ToM-1 Receiver dismisses any location that could be communicated by a shorter route as unlikely (“The Sender sent me message X, but could have reached her goal with the shorter message Y. Therefore, the additional states in message X are potentially my goal locations.”). Formally Simulated messages *m*_*sim*_ are the messages from starting location *A* to Sender’s goal location *G*_*s*_ and are shorter than *m*^*^:

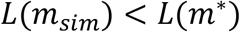

Next, the Receiver constructs 2 sets of unique states. One is unique states in the observed message: *S*_*m*_*(ob*s) *= U( m*^*^), and second is unique states in simulated shorter messages:

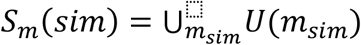

Where symbol big U denotes the union of all sets *U( m*_*sim*_) across each simulated message *m*_*sim*_. The Receiver identifies potential goal locations by calculating the set difference between these two sets:

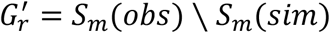

Finally, the Receiver randomly selects one state from 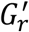:

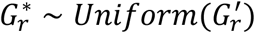

This **elimination strategy** is exemplified in Figure 6, where the ToM-1 Receiver refines his choices by considering the ToM-0 Sender’s perspective: he excludes 2 locations (figure 6c, yellow tiles) from his potential goals because the Sender would have opted for a shorter path to communicate those locations. The simulated shorter path is shown by the dashed line on Figure 6c, and the eliminated 2 states are the yellow-colored tiles. Thus, he chooses from the remaining two purple-colored locations with equal probability.

### ToM-2 players Sender

#### Mental model

ToM-2 Sender has the ToM-1 model of the Receiver, who himself mentalizes the Sender as the ToM-0 player. The ToM-2 Sender knows that the Receiver is employing the elimination strategy described above to narrow down the number of possible locations and improve his chances of identifying his goal location correctly.

#### Message selection

The ToM-2 Sender intentionally chooses a message that extends one step beyond the shortest path to the Receiver’s goal location. This additional step is designed to ensure that when the Receiver compares the observed path with potential shorter paths, the longer path clearly identifies the Receiver’s goal location. Figure 5c illustrates this approach, with a selected path (depicted in solid color) that purposefully diverges from the shorter alternatives (shown in lighter color). This strategy sacrifices reward (in the form of the shortest message) for clarity ensuring that the Receiver will identify his goal location without uncertainty.

Formally this process is described in the following way: the ToM-2 Sender model constructs 2 sets of messages: *M*_2,*direct*_, *M*_2,*pa*s*s*_. The first set *M*_2,*direct*_ consists of the messages that do not pass through the Receiver’s goal and go directly to the Sender’s goal. These are the shortest possible messages that avoid the Receiver’s goal. *M*_2,*direct*_ is defined as:

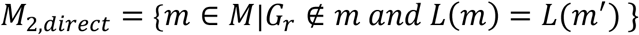

Second set is *M*_2,*pa*s*s*_ which consists of the messages that do pass through the Receiver’s goal. Each message in this set has one more unique state than those in *M*_2,*direct*_. Formally:

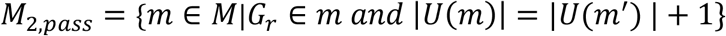

Finally, the Sender selects one message from *M*_2,*pa*s*s*_ randomly:

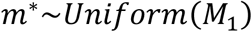

Meaning:

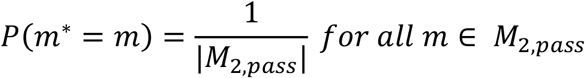

### Receiver

#### Mental model

ToM-2 Receiver has the ToM-1 model of the Sender, i.e. he knows that the Sender selects the path with the least possible locations.

#### Goal selection

The ToM-2 Receiver also uses an elimination strategy like the ToM-1 Receiver. But his mental model of the Sender is more efficient: as a ToM-2 Receiver, he anticipates that the ToM-1 Sender optimizes her message to have the least number of states. The ToM-2 Receiver’s elimination strategy is demonstrated in Figure 6d. Suppose, after observing the path in Figure 6a, the Receiver simulates two potential shorter messages the ToM-1 Sender could have sent. The first path is identical to the one simulated by the ToM-1 Receiver. With this simulated path, the Receiver excludes the 2 yellow tiles from possible goal locations (Figure 6d, left side grid) and considers 2 tiles as the potential goal locations (Figure 6d, left side grid, purple-colored tiles). The Receiver then simulates a second message (Figure 6d, right side grid) and by elimination strategy narrows down the candidate goal locations even further. He excludes the additional yellow tile from the candidate set, and now he knows with very high certainty that his goal location is the pink tile. Formally this process is described as follows.

ToM-2 Receiver observes *m*^*^ and identifies additional states included in the observed message but absent from messages with the fewer unique states, as these may be intentional indicators of the Receiver’s goal. He Simulates messages *m*_*sim*_ that include messages with fewer unique locations:

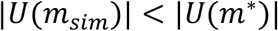

Next the Receiver constructs 2 sets of unique states. One is unique states in the observed message: *S*_*m*_*(ob*s) *= U( m*^*^), and second is unique states in simulated messages *m*_*sim*_:

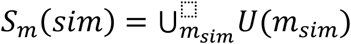

Where *m*_*sim*_ includes path with fewer unique states and symbol big U denotes the union of all sets *U( m*_*sim*_) across each simulated message *m*_*sim*_.

Finally, the Receiver identifies potential goal locations by calculating the set difference between these two sets:

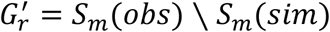

And selects one state from 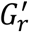 randomly:

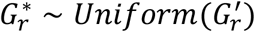

#### Variations in message selection across trial types and reasoning levels

It’s crucial to recognize that different levels of ToM reasoning lead to the selection of different message types. When applying these principles to indirect trials, ToM-0 yields what we term “pass-by” messages, while ToM-1 and ToM-2 results in the same “enter-exit” message (Figure 5b and 5c). However, in direct trials, the dynamics shift slightly. While the ToM-0 Sender still opts for the “pass-by” message type (as depicted in Figure 5d), at ToM-1, the Sender gravitates towards a “wiggly” message type (Message 2) contrary to the indirect trial Sender who selects “enter-exit” message (Figure 5b). To understand this strategic shift, consider Figure 5e, which compares two key messages: ‘Message 1’, a typical ‘enter-exit’ message, and ‘Message 2’, a typical ‘wiggly’ message. When the ToM-1 sender aims to minimize not only the number of moves (like the ToM-0 sender) but also the number of unique visited locations, message 2 appears to hold an advantage over message 1, namely that message 2 encompasses only 4 unique locations, whereas message 1 entails 5 unique locations. Consequently, the Sender opts for message 2 because it enhances the Receiver’s likelihood of randomly selecting the desired goal location from the visited locations.

### Surprise model

The core idea of the Surprise model in the Target Communication Game (TCG) is to reveal the Receiver’s target location by intentionally violating expected movement patterns. These patterns are set by probabilistic priors that predict the next moves and the likely path toward the Sender’s goal. By diverging from these patterns, the model identifies the most important parts of the message that the Receiver should focus on. To achieve this, the model calculates a surprise value for each possible action, choosing the one with the highest surprise. This process relies on accurately calculating these prior probabilities, which are divided into two types: movement priors and state priors

### Movement prior probabilities

This aspect of the model leverages the principles of movement kinetics to assign probabilities to each potential action. Movement physics implies that an object will likely continue its current path unless acted upon by an external force (figure 2e). Any deviation from this predicted path is thus deemed improbable and notably surprising. Thus, the probabilities assigned to each possible step are ordered as follows:

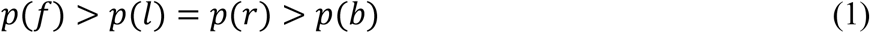

where *p(f*) is the probability of moving forward, *p(l*) and *p(r*) are the probabilities of moving left and right, respectively, and *p(b*) is the probability of moving backwards. These probabilities adjust dynamically with each movement, realigning based on the new direction of travel. In the Surprise Model, these probabilities are parameterized by two estimable parameters, expressed in the equations:

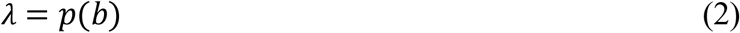

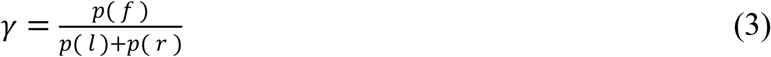

λ is generally set to be small between *0 .001* and *0 .3* whereas γ is adjusted within a range from 2 to 10. For instance, if γ *= 2*, then *p(f*) is twice as likely as the combined probability of a turn (*p(l*) + *p(r*)) and hence, *p(f*) *=* 2/3 and *p(l*) *= p(r*) *=* 1/6 (without adjusting for λ).

More generally, the probabilities for all four movement directions are derived from these two parameters as follows:

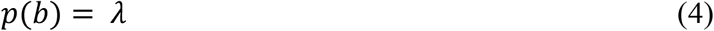

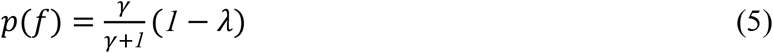

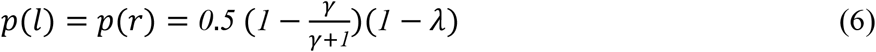

#### State prior probabilities

These priors are essential for establishing a baseline expectation of the Sender’s goal location (figure 2f). They define the probability of the state being at a given distance *d* from the Sender’s goal:

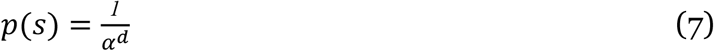

Here, p(s) represents the probability that a state is a distance *d* from the Sender’s goal. The parameter α, a positive estimable value, dictates how quickly these probabilities diminish as the distance from the Sender’s goal increases. While α stays the same across all trials, the specific point at which the state prior is the highest shifts with each new goal setup.

#### Integration of movement-state priors and calculating surprise

To calculate the final action probabilities *p(a)*, the model integrates movement and state priors through a multiplicative process and normalizes the results:

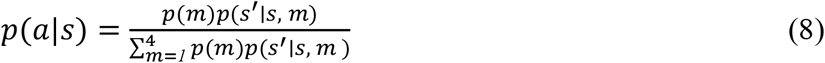

Here, *p(m)* denotes one of the four movement probabilities, and *p(s*′ ∣ *s, m* ) represents the state prior probability of state *s*′ after transitioning from state *s* with movement *m*. The Shannon surprise for each action is computed as:

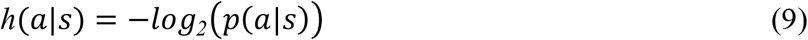

Where h(*a*|*s*) is a measure of information-theoretic surprise and shows how significantly an action deviates from expectations, with higher surprise values indicating greater deviation.

#### Rewards

If action choices were solely guided by Shannon surprise, the resulting path across the game board would wander inefficiently without directly progressing towards the Sender’s goal. In the TCG, rewards are conceptualized as the cost per step, incentivizing the Sender to quickly approach his target while maximizing the points preserved from the initial location. In this framework, the rewards for reaching the next state *s*′ from the current state *s* through action *a* are defined as the remaining points out of an initial endowment of 10 points for each goal configuration. After following the shortest path from the Receiver’s goal to the Sender’s goal. These rewards are then combined multiplicatively with the Shannon surprise calculated earlier to form an expected value:

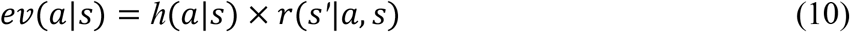

#### Two phases of planning

Crafting a message in the TCG involves two key steps: (1) Formulating a path to the Receiver’s goal that effectively communicates the location. This is achieved using the Surprise model, which selects actions based on how much they deviate from what is expected (measured by Shannon surprise). It also involves (2) subsequently navigating towards the Sender’s goal via the shortest possible route to conserve as much of the initial endowment as possible. During this second phase, the model does not utilize the Shannon surprise from the combined movement-state priors for choosing actions; instead, it follows the state priors directly towards the Sender’s goal. The expected value calculation for this phase is thus adjusted accordingly:

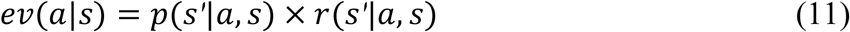

#### Planning horizon

In decision-making scenarios like the Target Communication Game (TCG), an agent can employ value iteration to strategize its forthcoming actions over a set period, known as the planning horizon^40^. This horizon specifies the number of steps the agent anticipates in advance. For instance, with a planning horizon of 2, the agent would calculate its moves from the current position through the next two steps. The objective of setting a planning horizon is to identify the sequence of actions that optimizes the total expected future rewards for the agent.

Within the framework of the Surprise model, the process involves creating a policy tree of all feasible actions and then assessing the expected values for each action in all branches of the policy tree while also accounting for possible future rewards. Specifically, the expected value of getting to the next state in the policy tree from the current state *s*_*i*_ using action *a*_*i*_ is the following:

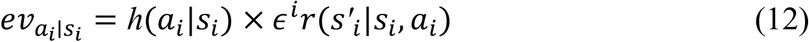

where h*(a*_*i*_|*s*_*i*_) is the surprise linked with the action probability of getting to the next state in the policy tree from the current state *s*_*i*_ using action *a*_*i*_. ϵ^*i*^ is the discount factor at the planning step *i*, and *r(s*′_*i*_|*s*_*i*_, *a*_*i*_) is the reward available in the next state *s*′ that is reached from the current state *s* via action *a* in the planning step *i* until the horizon is reached. After computing the expected values for each state-action pair in the tree, the model then aggregates these expected values across each path in the policy tree to determine the final expected values for all actions that are immediately available

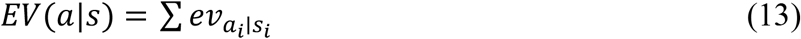

These expected values are subsequently processed using a softmax function to calculate the final probabilities for each action:

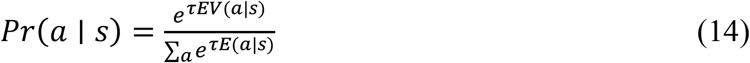

The softmax function computes the probability of selecting a specific action *a* in a given state *s*, factoring in the expected future rewards of each action. This calculation incorporates a temperature parameter τ that accounts for the randomness in the decision-making process.

## Data collection and analysis

### Sender’s data generation from computational models

To explore the dynamics of non-verbal communication in a Tacit Communication Game (TCG) we simulated Sender message generation through various computational models for selected 30 goal configurations. This set comprised 10 indirect goal configurations, which allowed for more straightforward communication, and 20 direct configurations, which required more complex inferential strategies. Sender messages were algorithmically generated based on distinct principles encapsulated by each model. For the Theory of Mind (ToM) models, messages were crafted considering the perspective-taking abilities at three different levels (ToM-0, ToM-1, ToM2). These levels reflect the degree to which a Sender can anticipate and respond to a Receiver’s cognitive state. In contrast, the Surprise model behavior was based on actions calculated to deviate from probabilistic expectations, thus signaling important information through the element of surprise.

Model-generated behavior was classified into discrete message types, which were defined by the pattern of the moves. These types included: Enter-exit, Wiggly and pass-by messages (figure 1d). These message types, each signifying a different communicative intent, were hypothesized to affect the Receiver’s ability to discern the intended target location correctly. Following the simulation of Sender behaviors, we analyzed the prevalence of each generated message type. The prevalence was expressed as the percentage that each message type constituted of the total messages produced by the computational models across all 30 goal configurations. Understanding the tendency of each model to favor certain communication strategies was essential for interpreting the subsequent accuracy and reaction time measures.

### Accuracy and reaction time measurement

The efficacy of each computational model was assessed by measuring the accuracy of human Receivers in correctly interpreting the transmitted message. This was done on two levels: the general accuracy of each model and the specific accuracy associated with each message type. Overall model accuracy was determined by the rate at which Receivers correctly inferred his own goal location. Additionally, we analyzed the accuracy rates for Enter-Exit, Wiggly, and Pass-By messages to discern the efficacy of each individual strategy in the communication process. Furthermore, reaction times, representing the speed of Receivers’ responses, were recorded to provide insight into the cognitive processing demands placed by each type of message.

We compared mean accuracies and reaction times in a one-way Analysis of Variance (ANOVA) with a predefined Type I error rate of p < 0.05 across the different message types and models using MATLAB. When the ANOVA revealed a significant difference, we employed post-hoc comparisons utilizing Tukey’s Honestly Significant Difference (HSD) test to examine significant differences between specific pairs of models.

## Acknowledgements

T.B., Y.Y., and J.G. are supported by the Collaborative Research Center TRR169 “Cross-modal Learning” funded by the German Research Foundation (DFG) and the National Science Foundation China (NSFC). J.G. is further supported by the Collaborative Research Center SFB1528 “Cognition of Interaction” funded by the German Research Foundation.

## Author contributions

All authors designed the research, T.B. and Y.Y. collected the data, J.G. designed and T.B. implemented the computational models, T.B. and Y.Y. analyzed the data. T.B. wrote the first draft of the manuscript; all authors contributed to the final version of the manuscript.

## Data availability statement

All behavioral and physiological data necessary to run the analysis is available on the authors’ GitHub at https://github.com/TatiaBu/Surprise_vs_ToM.

## Code availability statement

The code to run the analysis and reproduce the figures is archived and publicly available on GitHub at https://github.com/TatiaBu/Surprise_vs_ToM. The repository includes a README file with detailed instructions on how to set up the environment, install necessary dependencies, and execute the code.

## Notes

### Competing Interest Statement

The authors have declared no competing interest.

